# Defining the scope of the European Antimicrobial Resistance Surveillance network in Veterinary medicine (EARS-Vet): a bottom-up and One Health approach

**DOI:** 10.1101/2021.03.09.434124

**Authors:** Rodolphe Mader, on behalf of EU-JAMRAI, Clémence Bourély, Jean-Philippe Amat, Els M. Broens, Luca Busani, Bénédicte Callens, Paloma Crespo, Peter Damborg, Maria-Eleni Filippitzi, William Fitzgerald, Thomas Grönthal, Marisa Haenni, Annet Heuvelink, Jobke van Hout, Heike Kaspar, Cristina Munoz, Madelaine Norström, Karl Pedersen, Lucie Pokludova, Fabiana Dal Pozzo, Rosemarie Slowey, Anne Margrete Urdahl, Alkiviadis Vatopoulos, Christos Zafeiridis, Jean-Yves Madec

## Abstract

**Background:** Building the European Antimicrobial Resistance Surveillance network in Veterinary medicine (EARS-Vet) was proposed to strengthen the European One Health antimicrobial resistance (AMR) surveillance approach.

**Objectives:** The objectives were to (i) define the combinations of animal species, production types, age categories, bacterial species, specimens and antimicrobials to be monitored in EARS-Vet and to (ii) determine antimicrobial test panels able to cover most combinations.

**Methods:** The EARS-Vet scope was defined by consensus between 26 European experts. Decisions were guided by a survey of the combinations that are relevant and feasible to monitor in diseased animals in 13 European countries (bottom-up approach). Experts also considered the One Health approach and the need for EARS-Vet to complement existing European AMR monitoring systems coordinated by the European Centre for Disease Prevention and Control (ECDC) and the European Food Safety Authority (EFSA).

**Results:** EARS-Vet would monitor AMR in six animal species (cattle, swine, chicken (broiler and laying hen), turkey, cat and dog), for 11 bacterial species (*Escherichia coli*, *Klebsiella pneumoniae*, *Mannheimia haemolytica*, *Pasteurella multocida*, *Actinobacillus pleuropneumoniae*, *Staphylococcus aureus*, *Staphylococcus pseudintermedius*, *Staphylococcus hyicus*, *Streptococcus uberis*, *Streptococcus dysgalactiae* and *Streptococcus suis*). Relevant antimicrobials for their treatment were selected (e.g. tetracyclines) and complemented with antimicrobials of more specific public health interest (e.g. carbapenems). Three test panels of antimicrobials were proposed covering most EARS-Vet combinations of relevance for veterinary antimicrobial stewardship.

**Conclusions:** With this scope, EARS-Vet would enable to better address animal health in the strategy to mitigate AMR and better understand the multi-sectoral AMR epidemiology in Europe.

## Introduction

Antimicrobial resistance (AMR) cannot be tackled effectively without well-performing surveillance systems, covering the animal and human sectors.^1,2^ Currently, the European Centre for Disease Prevention and Control (ECDC) coordinates the European Antimicrobial Resistance Surveillance Network (EARS-Net), which monitors AMR in invasive bacterial isolates from patients with bloodstream infections or meningitis,^3^ and the European Food- and Waterborne Diseases and Zoonoses Network (FWD-Net), which monitors AMR in human *Salmonella* and *Campylobacter* infections.^4^ In the food sector, the European Food Safety Authority (EFSA) coordinates active monitoring of AMR in commensal and zoonotic bacteria from healthy food-producing animals (cattle, poultry, pigs) at slaughterhouse and food thereof, according to Directive 2003/99/CE and Commission Implementing Decision (EU) 2020/1729.^5^ As no European system currently monitors AMR in clinical isolates from diseased animals, data on AMR hotspots in specific animal infections are still lacking, although they are necessary to tailor strategies to rationalise antimicrobial usage in the animal sector.

In this context, the EU Joint Action on AMR and Healthcare Associated Infections (EU-JAMRAI), co-funded by the Third Health Programme of the EU, proposed to establish the European Antimicrobial Resistance Surveillance network in Veterinary medicine (EARS-Vet), to strengthen a One Health AMR surveillance approach in Europe.^6^ During EU-JAMRAI, it was agreed that EARS-Vet should work as a European network of national surveillance systems in diseased animals (similarly to EARS-Net) and complement and integrate with the AMR monitoring systems of the ECDC and EFSA. Its objectives would be to report on the AMR situation, follow AMR trends and detect emerging AMR in bacterial pathogens of animals in Europe. This information would contribute to:

- Advising policy makers on interventions to mitigate AMR.
- Monitoring the impact of European efforts to tackle AMR in the animal sector.
- Evaluating or revising marketing authorisations of antimicrobials.
- Supporting antimicrobial stewardship initiatives, especially the development of veterinary antimicrobial treatment guidelines.
- Generating missing epidemiological cut-off values and then clinical breakpoints for the interpretation of antimicrobial susceptibility testing (AST) results.
- Assessing the risk of AMR transmission between animals and humans via non-food related routes, e.g. by direct contact between humans and companion or food animals.
- Estimating the burden of AMR in animal health, e.g. attributable deaths caused by infections with antimicrobial-resistant bacteria.

A major step in the design of EARS-Vet was the definition of its surveillance scope, i.e. the combinations of animal species, production types, age categories, bacterial species, specimens (i.e. sample types) and antimicrobials to be monitored in EARS-Vet. The present study aims to describe these combinations and explain the bottom-up and One Health approach that was followed to determine them. This work also aims to propose a limited number of antimicrobial test panels that participating countries could use to cover most EARS-Vet combinations.

## Materials and methods

### 1. Collection of national veterinary AMR surveillance scopes

Thirteen countries shared their national veterinary AMR surveillance scope. This was done by completing an Excel template that recorded the combinations of animal species, production types, age categories, bacterial species, specimens and antimicrobials monitored by countries with a national surveillance system (the Czech Republic, Denmark, Estonia, Finland, France, Germany, Ireland, the Netherlands, Norway and Sweden), or considered as future monitoring targets in countries that were in the process of establishing a surveillance system (Spain), or planning to build their system (Belgium and Greece), at the time of study. The national surveillance scopes of Belgium, Greece and Spain were defined *via* the consultation of relevant national experts, who were advised to follow the recommendations of the World Organisation for Animal Health (OIE) in this regard.^7^

### 2. Identification of the feasibility to widen national veterinary AMR surveillance scopes

Representatives of the 13 countries sharing their national veterinary AMR surveillance scope were also questioned about the feasibility to monitor AMR in clinical animal isolates for seven of the eight bacterial species monitored by EARS-Net (*Escherichia coli, Pseudomonas aeruginosa, Klebsiella pneumoniae, Acinetobacter baumannii, Staphylococcus aureus, Enterococcus faecalis* and *Enterococcus faecium*) among the animal species already included in their national surveillance scope. *Streptococcus pneumoniae*, also monitored in EARS-Net, was not included in this survey as it is primarily a human pathogen. For each combination of animal species / bacterial species, countries had the choice between three answers: (i) the monitoring is feasible and the combination is already included in the national scope (“feasible and included”), (ii) the monitoring is feasible but the combination is not included in the national scope (“feasible but not included”) or (iii) the monitoring is “currently not feasible”. Justifications were required in case of answering (ii) or (iii).

### 3. Expert consensus on the EARS-Vet surveillance scope

The combinations to be included in the EARS-Vet surveillance scope were defined by consensus among 26 experts (veterinary and human microbiologists, veterinary epidemiologists and Ministry representatives) from 14 European countries (the 13 countries that shared their national scope and Italy), who met at six teleconferences in 2020. Expert discussions were guided by a preliminary analysis of the data collected, and they considered the need for EARS-Vet to complement existing monitoring systems coordinated by the ECDC and EFSA. The decision process followed this stepwise approach:

1. Determination of the animal species.
2. Determination of the bacterial species within each selected animal species.
3. Determination of accepted clinical specimens for each combination of animal species / bacterial species.
4. Determination of surveillance stratification per production type and / or age category for each combination of animal species / bacterial species / specimen.
5. Determination of the antimicrobials for each combination of animal species / bacterial species / specimen / production type / age category.

## Results

### 1. Determination of the animal species

Figure 1 shows the animal species included in the 13 national veterinary AMR surveillance scopes. All countries reported major food-producing animals (cattle, swine, chicken or turkey), nine included cats and dogs, and four included horses. The expert panel decided to monitor in EARS-Vet the six animal species most frequently included in national surveillance scopes: cattle, swine, chickens, cats, dogs, and turkeys.

**Figure 1:**
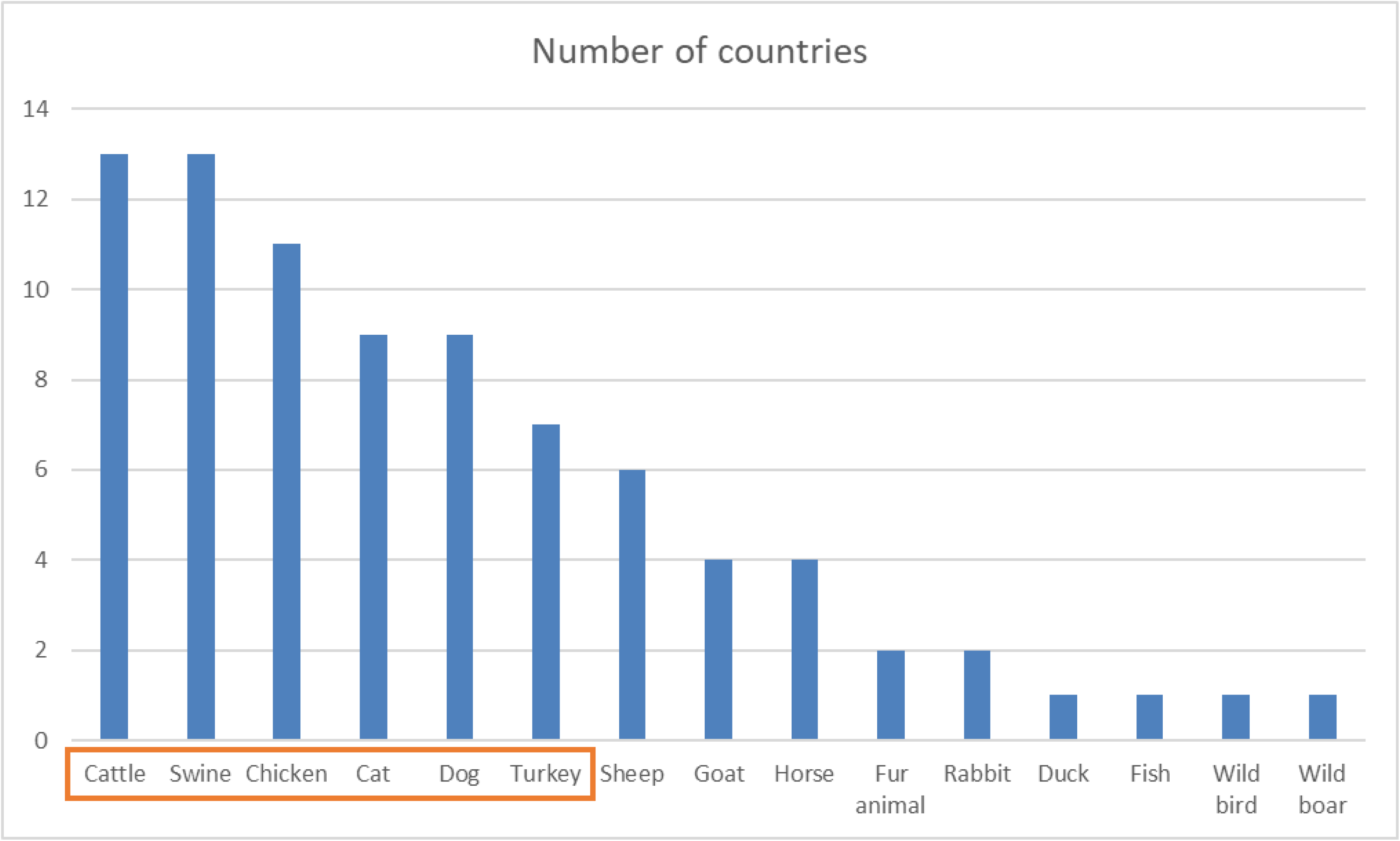
Distribution of animal species selected in the national surveillance scopes of 13 European countries

### 2. Determination of the bacterial species within each selected animal species

Figure 2 shows the bacterial species included in the 13 national veterinary AMR surveillance scopes for each of the six chosen animal species. The expert group decided to include the most frequently reported bacterial species within each animal species in the EARS-Vet scope. No threshold defined the selection of bacterial species, but in practice, none was selected if included by fewer than three countries. *Salmonella* spp. was not selected, considering that many efforts are already in place in Europe to monitor its AMR in the animal sector (EFSA monitoring). *E. coli* is the only bacterial species included for all six selected animal species.

**Figure 2:**
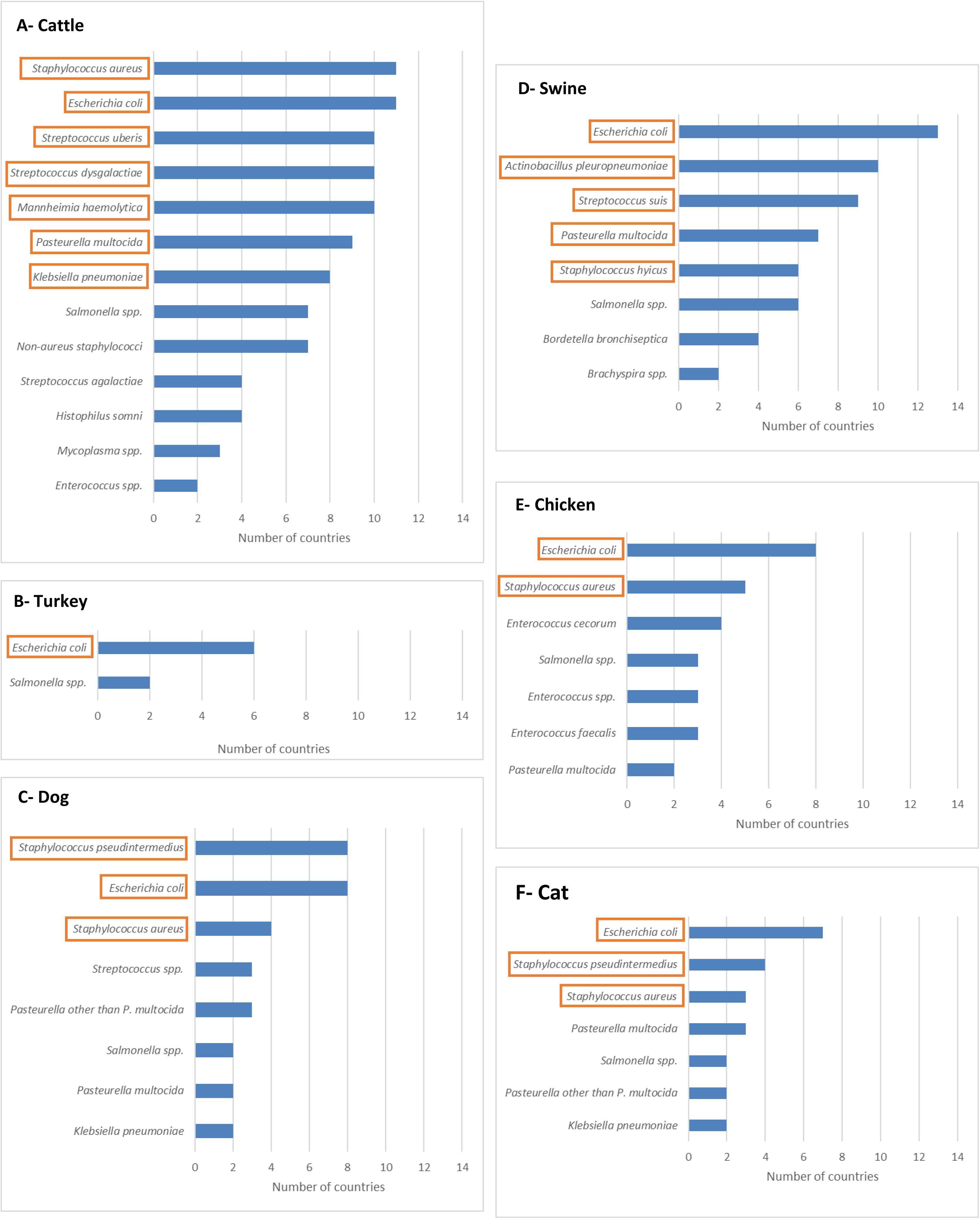
Distribution of bacterial species selected in the national scopes of 13 European countries per cattle, swine, turkey, chickens, cats and dogs

Table 1 shows the feasibility of monitoring also AMR in clinical animal isolates representing the seven bacterial species of public health interest (monitored in EARS-Net). The combinations that most countries found feasible and already had in their surveillance scope were *E. coli* in cattle and swine, and *S. aureus* in cattle, whereas no country included *A. baumannii*. Combinations of bacteria and animal species that were categorised “feasible but not included” were assessed in order to evaluate whether countries could expand their surveillance systems as part of the EARS-Vet programme. For only 16 (out of 42) animal species / bacterial species combinations the answer “feasible but not included” was given. However the number of countries never exceeded three, except for the combination dog / *E. faecalis*. The most frequent reason for such an answer was that the bacterium was not a surveillance priority. The most common reason for reporting “currently not feasible” for an EARS-Net bacterium was that it was too rarely obtained from animal clinical specimens (countries used thresholds ranging from 10 to 100 isolates minimum per year). Finally, the expert group concluded that the EARS-Vet scope should not include more bacterial species than those initially selected, based on national scopes.

**Table 1:**
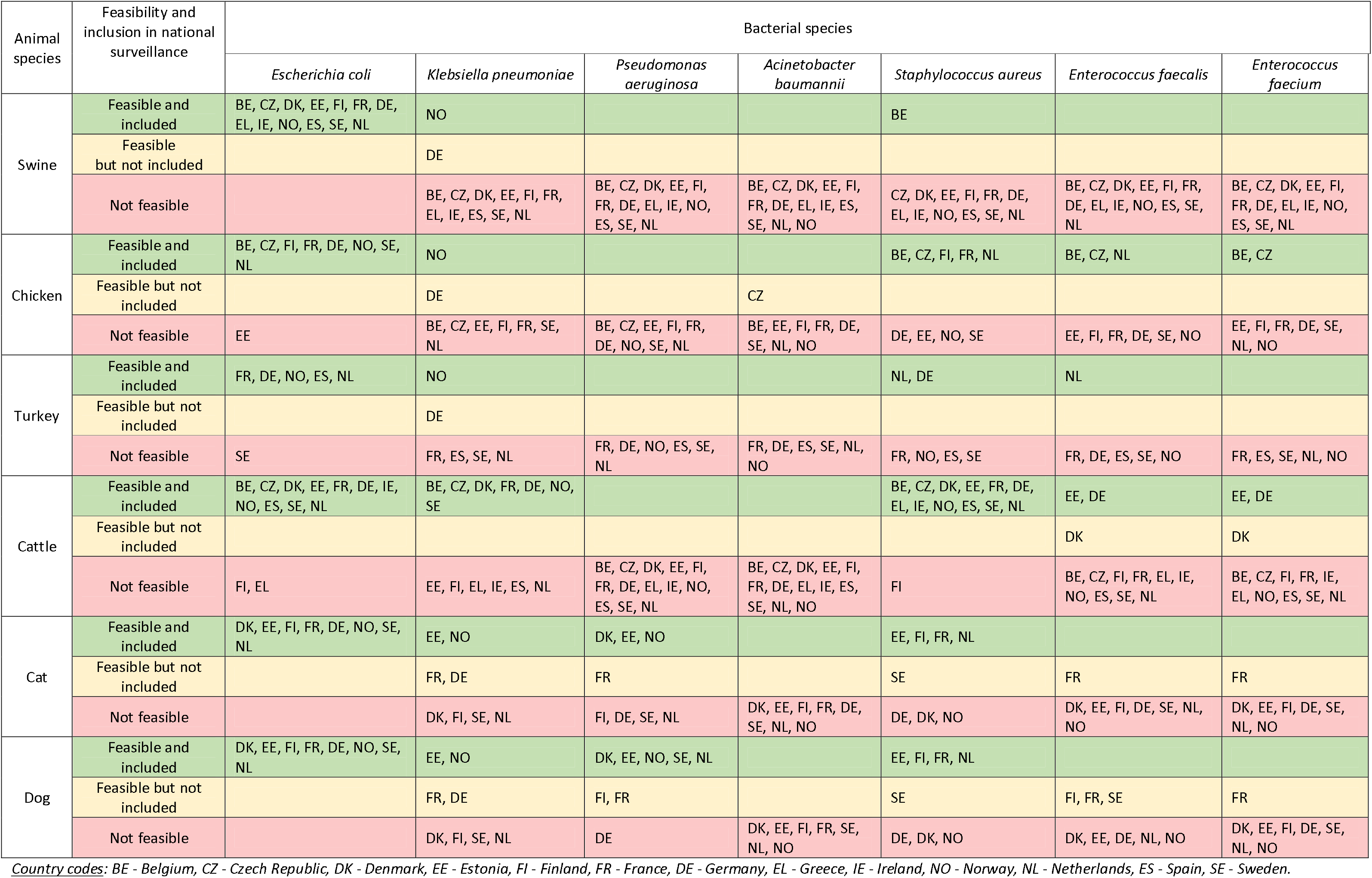
Countries’ self-assessment of the feasibility to monitor AMR in clinical animal isolates for seven bacterial species of public health interest (monitored in EARS-Net) in the animal species included in EARS-Vet

### 3. Determination of accepted specimens for each animal species / bacterial species combination

Table 2 summarises decisions taken by the expert group on the specimens that would be accepted in EARS-Vet. The surveillance of AMR would be stratified per specimen only for *E. coli* in cattle, which would be monitored separately in milk samples (in case of mastitis), in inner organs or blood (in case of septicaemia) and in faeces / intestinal content (in case of diarrhoea). In swine, it was decided to monitor *E. coli* in faeces / intestinal content only, as national surveillance systems rarely monitor it in blood or inner organs. When it comes to *E. coli* from swine faecal / intestinal samples, most countries monitor AMR only for virulent isolates or stratify their surveillance between virulent and non-virulent isolates, identified by PCR, serotyping, haemolytic profile, or a combination of them. Thus, the expert group decided to collect information on the haemolytic profile, serotypes and virulence factors of swine *E. coli* to stratify AMR data based on typing information.

**Table 2:**
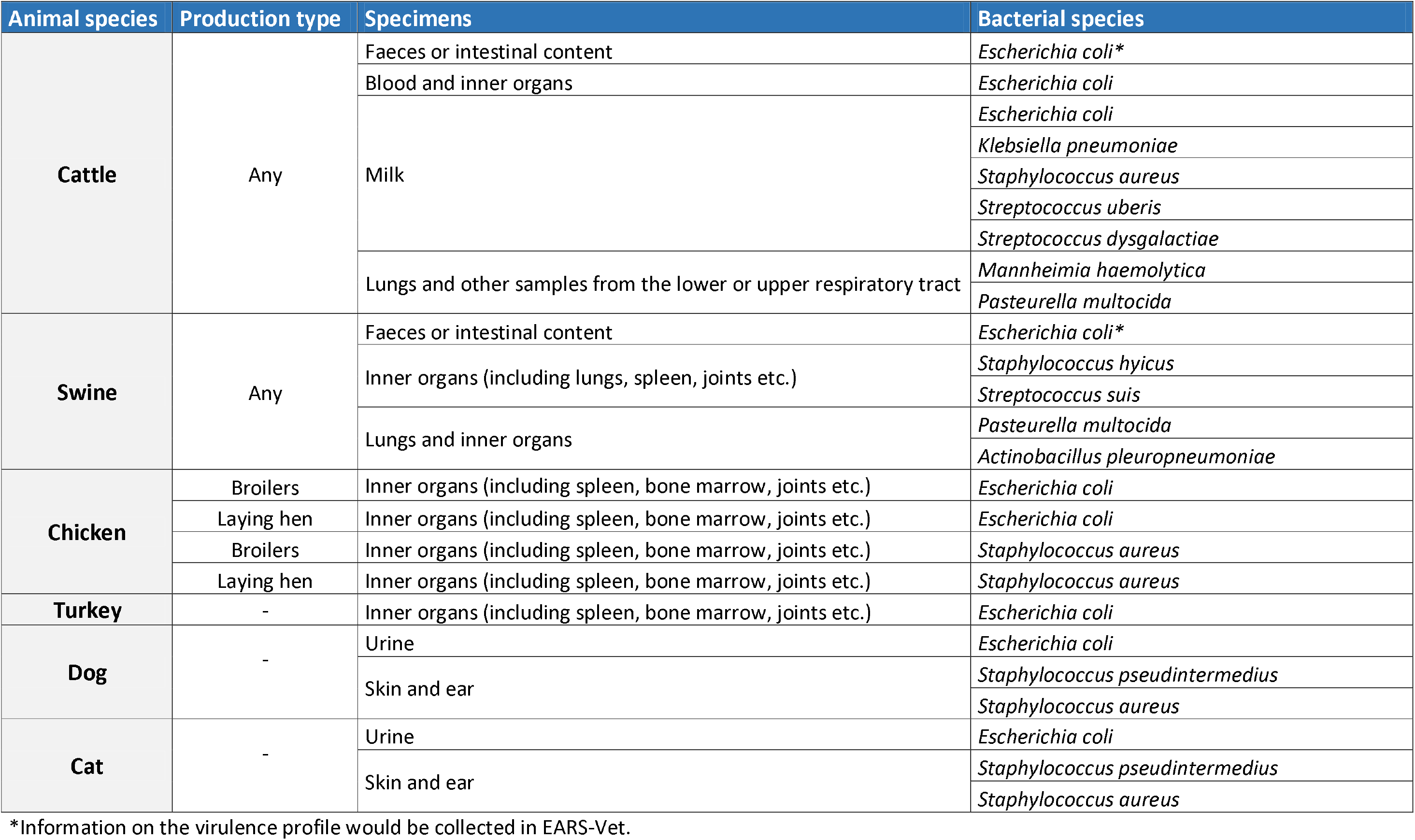
Animal species, production types, specimens and bacterial species to be covered by the European Antimicrobial Resistance Surveillance network in Veterinary medicine (EARS-Vet)

### 4. Determination of stratification per production type and age category for each animal species / bacterial species / specimen combination

Some participating countries stratify their national surveillance system per production type and age category. For instance, in the Czech Republic, the surveillance of pathogens in swine is stratified between pre-weaning piglets, post-weaning piglets, fattening pigs and sows. However, this is not systematic across national surveillance systems. Experts pointed out that this information was often missing in their national databases, except for chicken, where broiler and laying hen are most often distinguished. Therefore, EARS-Vet would only collect information on production type for chicken and no information would be collected on age category (Table 2).

### 5. Determination of antimicrobials for each animal species / bacterial species / specimen / production type combination

Different countries often monitor different antimicrobial agents for a given combination of animal species / bacterial species / specimen / production type. Therefore, the expert group decided that EARS-Vet should monitor AMR for:

- antimicrobial classes (e.g. tetracyclines), when it is possible to define class representatives (e.g. doxycycline).
- individual antimicrobial agent (e.g. neomycin) when class representatives cannot be defined (e.g. aminoglycosides).
- specific resistance phenotypes, e.g. methicillin resistance in staphylococci.

When it comes to an entire antimicrobial class or a specific resistance phenotype, EARS-Vet would accept and collate AMR data that correspond to different antimicrobial agents, as in EARS-Net.^3^

Figures 3 to 8 show which antimicrobial categories would be included for each combination of animal species and bacterial species. In general, the ones most frequently included in national surveillance scopes were selected for the EARS-Vet scope, with a few notable exceptions:

- Cephalosporins and penicillins + beta-lactamase inhibitors were not selected for *Pasteurella multocida, Mannheimia haemolytica* and *Actinobacillus pleuropneumoniae*, as resistance to beta-lactams is not common in Europe and penicillins alone are already included.^3^
- Third-generation cephalosporins were not selected for staphylococci, as *in vitro* susceptibility testing of these agents is less reliable than other antimicrobials testing resistance to cephalosporins indirectly (i.e. those used for monitoring methicillin resistance).
- Phenicols were not selected when they were not commonly used to treat the condition in question, although frequently included in national scopes for several combinations.
- Sulphonamides and trimethoprim were not included as separate classes in swine / *E. coli*, as they are mostly used as fixed combination in clinical practice for *E. coli* associated infections.
- Tildipirosin was not included for respiratory pathogens, as two other macrolides were already included in the EARS-Vet scope: tulathromycin and tilmicosin.
- Fusidic acid was not included for *S. aureus* and *Staphylococcus pseudintermedius* in cats and dogs, as this drug and its derivatives are only used for topical treatment in veterinary medicine. In that regard, little evidence exists between AST results and clinical efficacy when it comes to topical agents.

**Figure 3:**
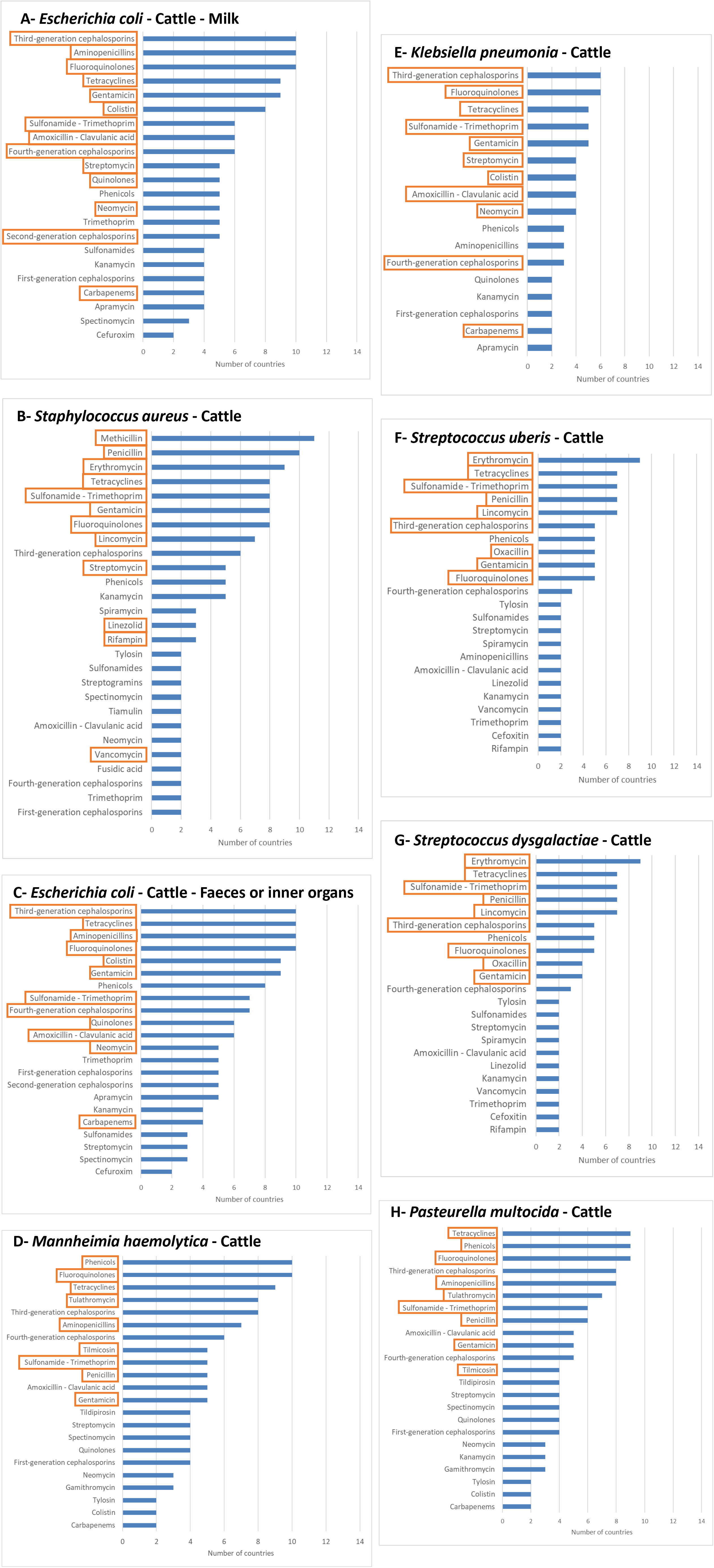
Distribution of antimicrobial categories selected in national surveillance scopes for *E. coli, K. pneumoniae, S. aureus, S. uberis, S. dysgalactiae, P. multocida* and *M. haemolytica* isolated from cattle

**Figure 4:**
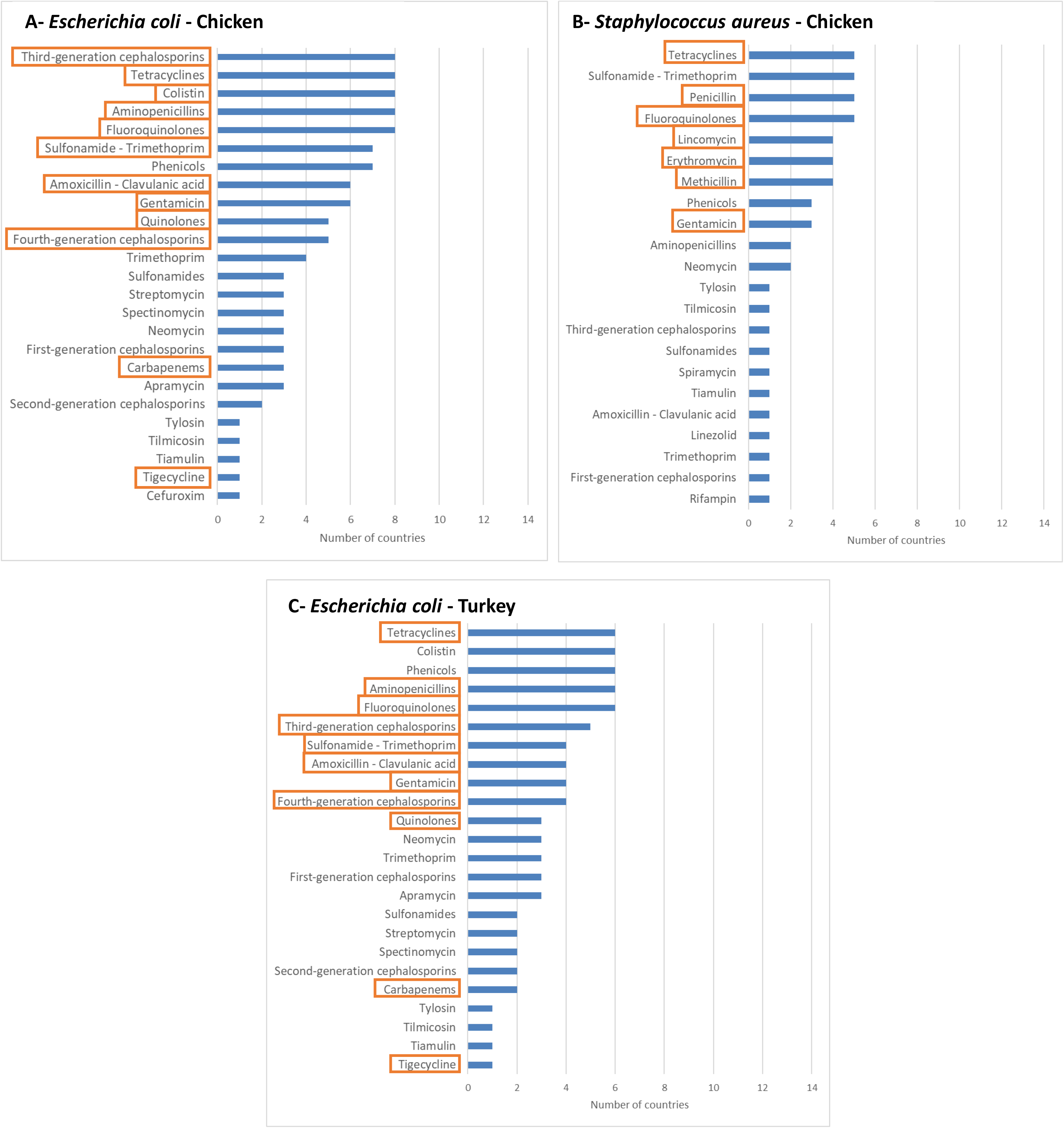
Distribution of antimicrobial categories selected in national surveillance scopes for *E. coli* and *S. aureus* isolated from chickens and *E. coli* isolated from turkeys

**Figure 5:**
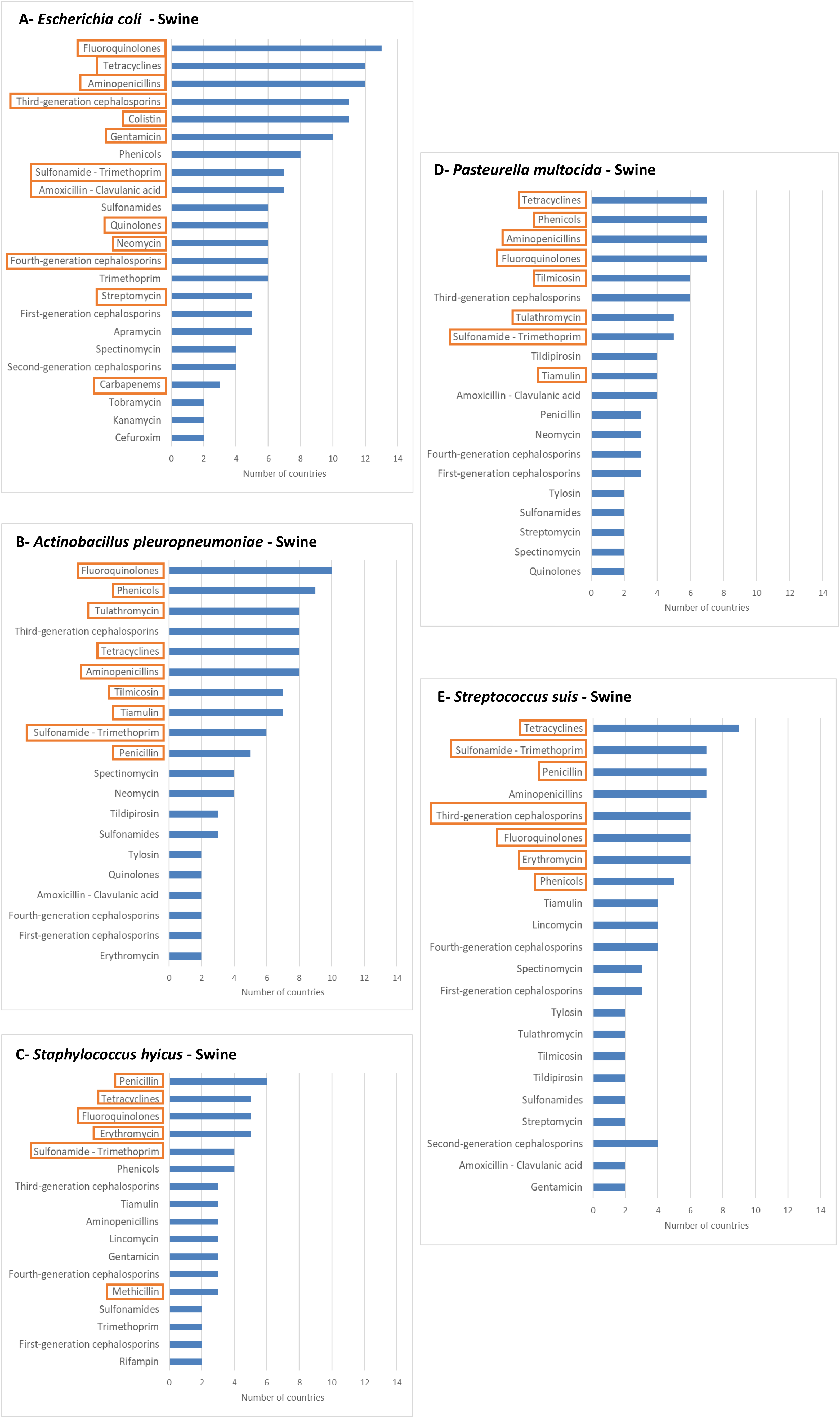
Distribution of antimicrobial categories selected in national surveillance scopes for *E. coli, S. suis, S. hyicus, A. pleuropneumoniae* and *P. multocida* isolated from swine

**Figure 6:**
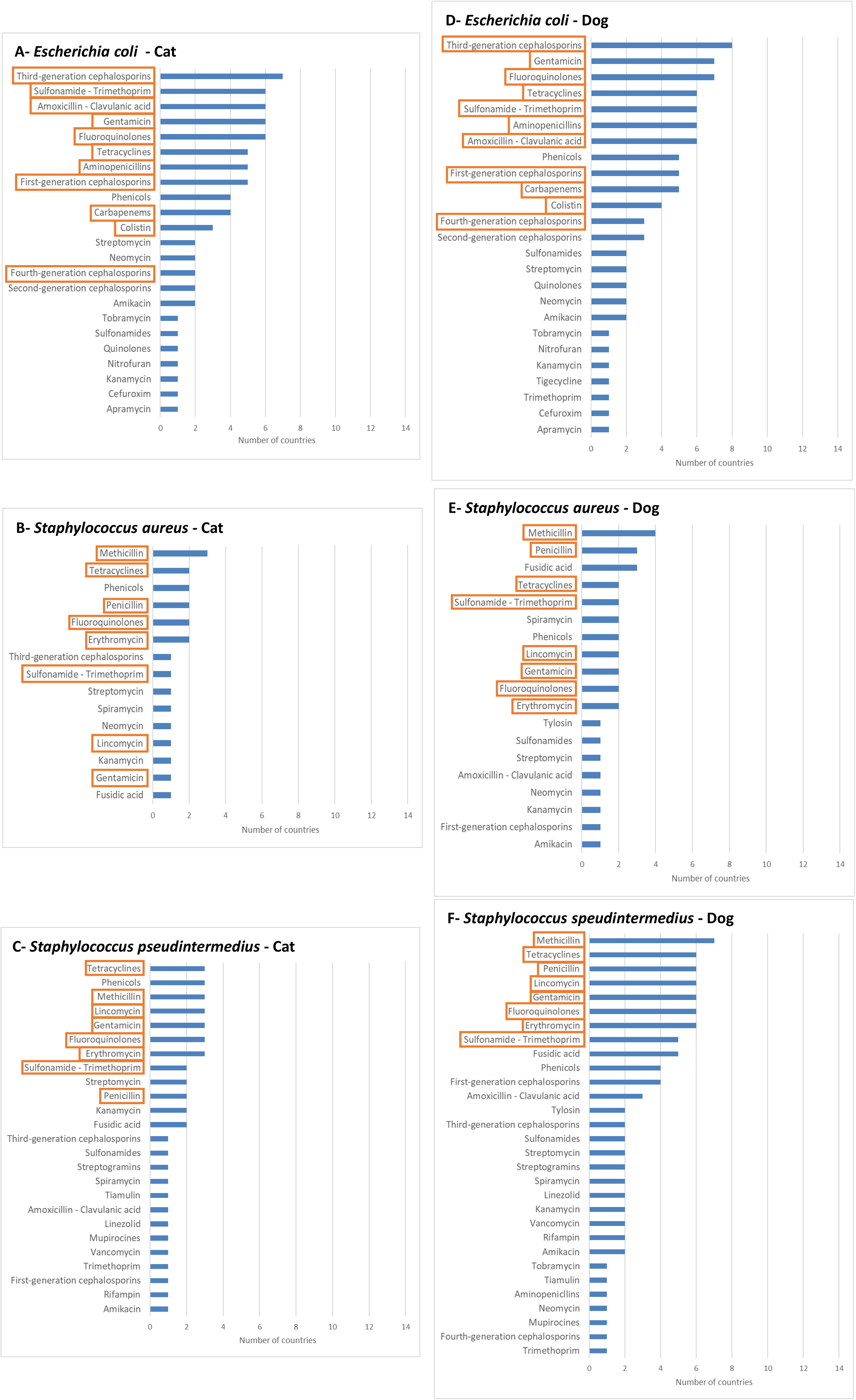
Distribution of antimicrobial categories selected in national surveillance scopes for *E. coli, S. aureus* and *S. pseudintermedius* isolated from cats and dogs.

For bacterial species included in both EARS-Vet and EARS-Net (no common bacterial species between EARS-Vet and the AMR monitoring component of FWD-Net), all AST results to antimicrobial classes that are important from a public health perspective and monitored by EARS-Net would also be included, even though they were not frequently included in national scopes. Thus, for *E. coli* and *K. pneumoniae*, EARS-Vet would also collect AST results for piperacillin-tazobactam, carbapenems and tigecycline and, for *S. aureus*, vancomycin, rifampin, linezolid and daptomycin.

A list of antimicrobial agents was determined per each antimicrobial class or specific resistance phenotype selected per combination. For each combination, countries would only report AMR data for the compounds on the lists. The lists aim to maximise country participation and minimise the diversity of accepted antimicrobial agents by EARS-Vet. In the case of isolated *E. coli* from cats, countries reported in their surveillance scopes: enrofloxacin, ciprofloxacin, marbofloxacin, danofloxacin among fluoroquinolones. However, all countries reporting marbofloxacin and/or danofloxacin also reported enrofloxacin, whereas some countries reported only enrofloxacin or only ciprofloxacin. Thus, the list for *E. coli* from cats comprises only enrofloxacin and ciprofloxacin. Appendix 1 *(=Tables 3 to 6)* summarises the EARS-Vet scope at antimicrobial category and compound levels.

**Table 3:**
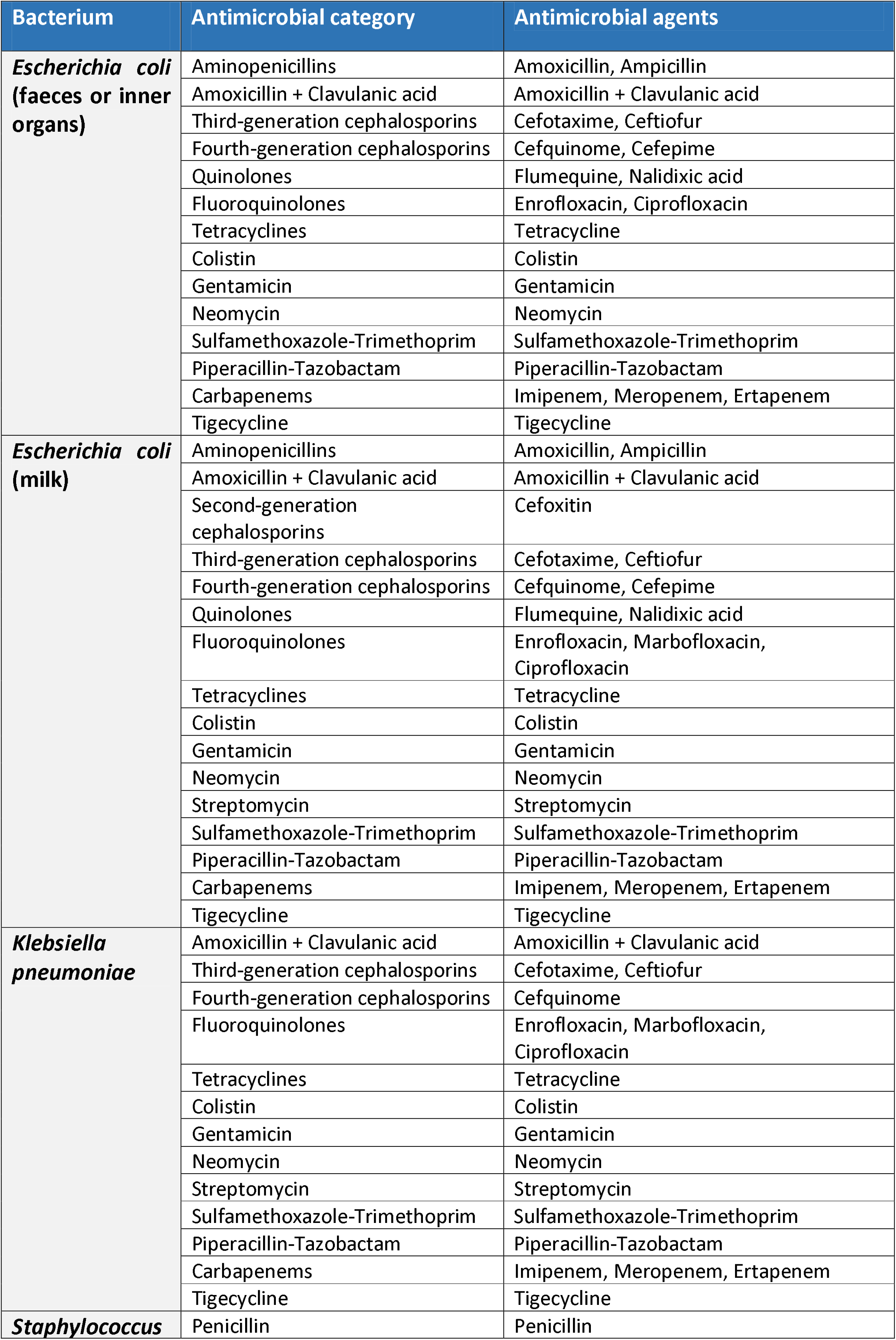
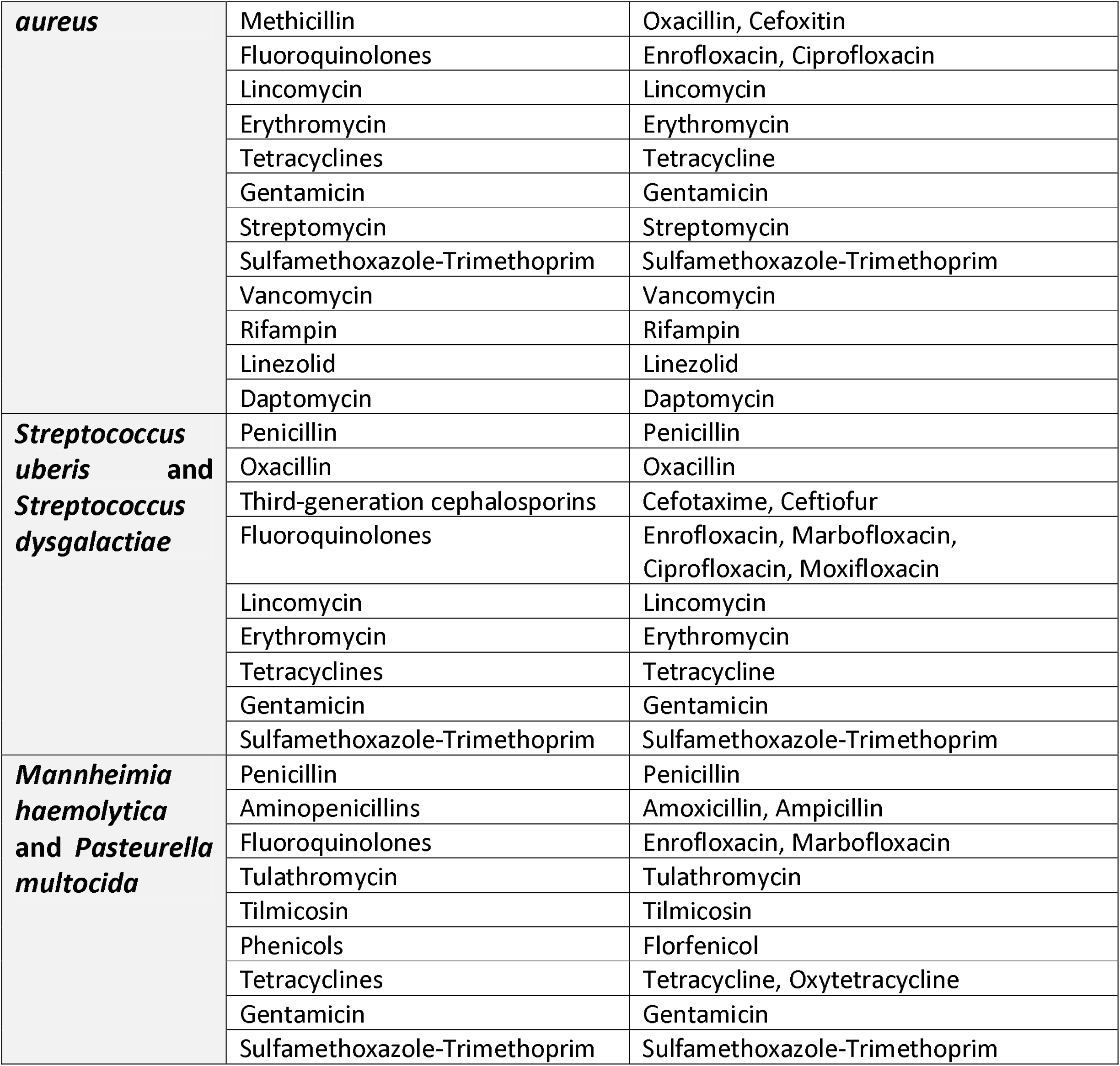
Bacterium-antimicrobial combinations included in the EARS-Vet scope for cattle

**Table 4:** Bacterium-antimicrobial combinations included in the EARS-Vet scope for swine

| Bacterium | Antimicrobial category | Antimicrobial agents |
| --- | --- | --- |
| ***Escherichia coli*** | Aminopenicillins | Amoxicillin, Ampicillin |
|  | Amoxicillin + Clavulanic acid | Amoxicillin + Clavulanic acid |
|  | Third-generation cephalosporins | Cefotaxime, Ceftiofur |
|  | Fourth-generation cephalosporins | Cefquinome, Cefepime |
|  | Quinolones | Flumequine, Nalidixic acid |
|  | Fluoroquinolones | Enrofloxacin, Ciprofloxacin |
|  | Tetracyclines | Tetracycline |
|  | Colistin | Colistin |
|  | Gentamicin | Gentamicin |
|  | Neomycin | Neomycin |
|  | Streptomycin | Streptomycin |
|  | Sulfamethoxazole-Trimethoprim | Sulfamethoxazole-Trimethoprim |
|  | Piperacillin-Tazobactam | Piperacillin-Tazobactam |
|  | Carbapenems | Imipenem, Meropenem, Ertapenem |
|  | Tigecycline | Tigecycline |
| ***Staphylococcus hyicus*** | Penicillin | Penicillin |
|  | Methicillin | Oxacillin, Cefoxitin |
|  | Fluoroquinolones | Enrofloxacin, Ciprofloxacin |
|  | Erythromycin | Erythromycin |
|  | Tetracyclines | Tetracycline |
|  | Sulfamethoxazole-Trimethoprim | Sulfamethoxazole-Trimethoprim |
| ***Streptococcus suis*** | Penicillin | Penicillin |
|  | Third-generation cephalosporins | Ceftiofur |
|  | Fluoroquinolones | Enrofloxacin, Ciprofloxacin |
|  | Erythromycin | Erythromycin |
|  | Phenicol | Florfenicol, Chloramphenicol |
|  | Tetracyclines | Tetracycline |
|  | Sulfamethoxazole-Trimethoprim | Sulfamethoxazole-Trimethoprim |
| ***Pasteurella multocida* and *Actinobacillus pleuropneumoniae*** | Penicillin (only for *A. pleuropneumoniae*) | Penicillin |
|  | Aminopenicillins | Amoxicillin, Ampicillin |
|  | Fluoroquinolones | Enrofloxacin, Marbofloxacin, Ciprofloxacin |
|  | Tulathromycin | Tulathromycin |
|  | Tilmicosin | Tilmicosin |
|  | Phenicol | Florfenicol, Chloramphenicol |
|  | Tetracyclines | Tetracycline, Oxytetracycline, Doxycycline |
|  | Sulfamethoxazole-Trimethoprim | Sulfamethoxazole-Trimethoprim |
|  | Tiamulin | Tiamulin |

**Table 5:** Bacterium-antimicrobial combinations included in the EARS-Vet scope for chicken and turkey

| Bacterium | Antimicrobial category | Antimicrobial agents |
| --- | --- | --- |
| ***Escherichia coli* (chicken and turkey)** | Aminopenicillins | Amoxicillin, Ampicillin |
|  | Amoxicillin + Clavulanic acid | Amoxicillin + Clavulanic acid |
|  | Third-generation cephalosporins | Cefotaxime, Ceftiofur |
|  | Fourth-generation cephalosporins | Cefquinome, Cefepime |
|  | Quinolones | Flumequine, Nalidixic acid |
|  | Fluoroquinolones | Enrofloxacin, Ciprofloxacin |
|  | Tetracyclines | Tetracycline, Doxycycline |
|  | Colistin | Colistin |
|  | Gentamicin | Gentamicin |
|  | Sulfamethoxazole-Trimethoprim | Sulfamethoxazole-Trimethoprim |
|  | Piperacillin-Tazobactam | Piperacillin-Tazobactam |
|  | Carbapenems | Imipenem, Meropenem, Ertapenem |
|  | Tigecycline | Tigecycline |
|  | Penicillin | Penicillin |
| ***Staphylococcus aureus* (only for chicken)** | Methicillin | Oxacillin, Cefoxitin |
|  | Fluoroquinolones | Enrofloxacin, Ciprofloxacin |
|  | Lincomycin | Lincomycin |
|  | Erythromycin | Erythromycin |
|  | Tetracyclines | Tetracycline, Doxycycline |
|  | Gentamicin | Gentamicin |
|  | Sulfamethoxazole-Trimethoprim | Sulfamethoxazole-Trimethoprim |
|  | Vancomycin | Vancomycin |
|  | Rifampin | Rifampin |
|  | Linezolid | Linezolid |
|  | Daptomycin | Daptomycin |

**Table 6:**
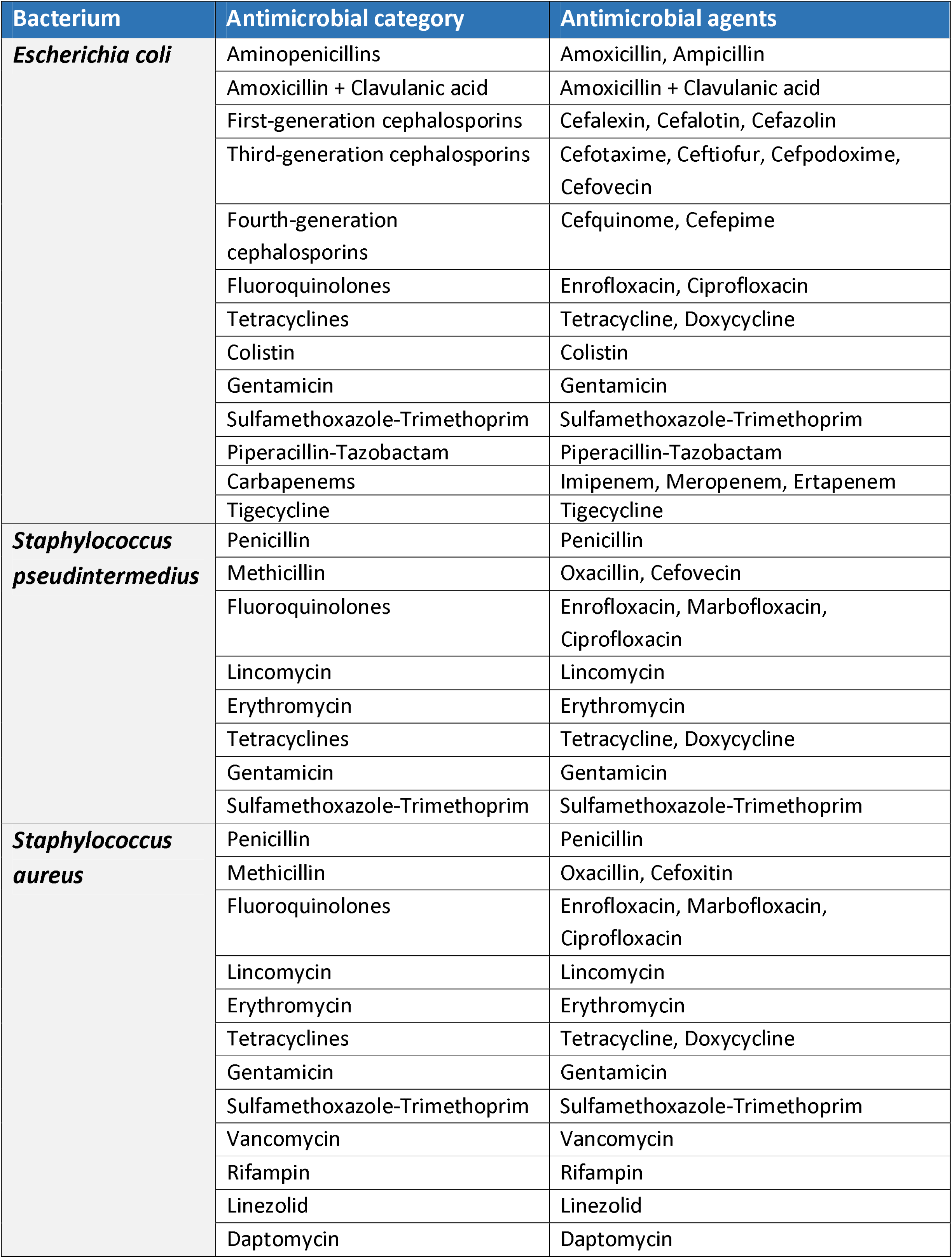
Bacterium-antimicrobial combinations included in the EARS-Vet scope for cats and dogs

Finally, by selecting the most frequently included antimicrobial compound for each combination, three test panels of antimicrobials were proposed to cover nearly all combinations of the EARS-Vet surveillance scope:

- Panel A for *E. coli* and *K. pneumoniae*: ampicillin, amoxicillin + clavulanic acid, cefotaxime or ceftiofur (cefotaxime being preferred for the detection of extended spectrum beta-lactamase producing *Enterobacterales*), cefquinome, nalidixic acid or flumequine, enrofloxacin, tetracycline, colistin, gentamicin and sulfamethoxazole - trimethoprim.
- Panel B for *P. multocida, M. haemolytica* and *A. pleuropneumoniae*: penicillin or ampicillin, enrofloxacin, tulathromycin, tilmicosin, florfenicol, tetracycline and sulfamethoxazole - trimethoprim.
- Panel C for *S. aureus, S. pseudintermedius, Streptococcus suis, Streptococcus dysgalactiae* and *Streptococcus uberis*: penicillin, oxacillin (to detect methicillin-resistant *S. pseudintermedius*), cefoxitin (to detect methicillin-resistant *S. aureus*), enrofloxacin, lincomycin, erythromycin, tetracycline, gentamicin and sulfamethoxazole - trimethoprim.

Countries may of course supplement these test panels with antimicrobials of more local clinical relevance.

## Discussion

In this work, consensus among experts was reached on the combinations of animal species, bacterial species, specimens, production types and antimicrobials to be monitored in EARS-Vet. Although with a primary focus on animals, strong efforts were made to define a scope that is relevant from a One Health perspective and that would enable EARS-Vet to complement and integrate with EARS-Net and the EFSA monitoring. Moreover, this work allowed the formation of three test panels which cover most combinations of the EARS-Vet scope and could be useful resource for countries building their surveillance system.

This work was based on 13 national surveillance scopes. Such a bottom-up approach proved particularly helpful to guide expert discussions, by providing information about what is relevant and feasible to monitor in a large part of Europe. Most of the ten countries with a national surveillance system in place are Northern and Central European countries, but the inclusion in this work of two Southern European countries, Spain and Greece, improved European representativeness. Collecting the national scope of countries building or planning to build their system also enabled to anticipate a future expansion of the network. However, they might not have the experience to assess accurately their capacity to monitor some combinations. For countries with a system in place, a strength of our approach has been direct interaction with national coordinators, rather than solely relying on AMR surveillance reports as a source of information. It helped locate and share non-published data in some countries. It revealed that reported antimicrobials are not always those actually tested, e.g. ciprofloxacin may be tested, but enrofloxacin is reported, as ciprofloxacin is not authorized for animal use. It enabled us to better understand what epidemiological data are collected by surveillance systems, e.g. when AMR data are reported for poultry and not per poultry species. Finally, some countries need to collect certain data over several years before being able to report them nationally. With our interactive approach with coordinators, we could overcome these challenges and consider more recent combinations being monitored, for example when test panels have changed since their last surveillance report publication.

The expert group decided to include six animal species in the EARS-Vet scope, covering both food-producing and companion animals. The three countries that were either building or planning to build their system (Spain, Belgium and Greece) did not include companion animals in their scope, as focussing on food-producing animals was the priority at their stage. Notably, the EARS-Vet scope includes all food-producing animal species covered by the EFSA monitoring of AMR in commensal and zoonotic bacteria and endeavours to fill a current surveillance gap in Europe with the inclusion of dogs and cats.

Major bacterial pathogens, considered as important drivers for veterinary antimicrobial usage in Europe, were selected for each animal species. Three of them are also of strong public health relevance (*E. coli*, *S. aureus* and *K. pneumoniae*) and could act as surveillance indicators across animal species and European AMR monitoring systems. Previous studies have shown different proportions and evolutions of AMR between commensal and pathogenic *E. coli* in food-producing animals,^8,9^ and emphasise the importance of monitoring AMR in both *E. coli* populations through EFSA and EARS-Vet, respectively.

Different specimens were selected in the EARS-Vet scope, depending on the animal species / bacterial species combination. Invasive samples, if they are not contaminated, ensure that isolated bacteria are the cause of infection. However, it can be challenging or impossible to identify the cause of infection for samples taken in sites with a commensal flora or when there is a high risk of contamination. This is the case, for example, where *E. coli* is isolated of faecal samples. Although laboratories usually investigate the serotypes or virulence factors of *E. coli* from piglets, it is not commonly performed in *E. coli* isolates from calves. This would introduce a source of bias in the surveillance of EARS-Vet to be considered when interpreting results. By comparison with the human sector, EARS-Net includes only invasive samples (from blood and cerebrospinal fluid), but this is not the case of the Global Antimicrobial Resistance Surveillance System (GLASS), coordinated by the World Health Organization, which also includes for instance stool samples to perform surveillance on *Salmonella* spp. and *Shigella* spp.^10^ Although stratification per production type would only be done in chickens (between broilers and laying hens), and no stratification would be done by age category, several combinations could be associated to a specific production type or age category, without collecting this information. For example, mastitis pathogens would almost exclusively originate from adult dairy cows and *E. coli* from swine faecal samples typically from piglets with neonatal or post-weaning diarrhoea.

Regarding antimicrobials, our bottom-up and One Health approach led to the selection of antimicrobials of both animal and human relevance, covering all categories of the Antimicrobial Advice *ad hoc* Expert Group (AMEG) of the European Medicines Agency (EMA). These categories were defined based on their potential consequences to public health (due to increased AMR) when used in animals, but also considering the need to use them in veterinary medicine.^11^ In a perspective of integration, for the three common bacterial species of EARS-Net and EARS-Vet, an important decision has been to include the antimicrobials monitored by EARS-Net in the EARS-Vet scope too. However, antimicrobial groupings may vary between these surveillance systems, as tested antimicrobial compounds often differ in veterinary and human medicine. For instance, EARS-Net collates resistance data to ciprofloxacin, levofloxacin and ofloxacin to monitor fluoroquinolone resistance in *E. coli*,^3^ whereas EARS-Vet would accept AST results for enrofloxacin, marbofloxacin and ciprofloxacin for *E. coli* from cattle mastitis. In contrast to EARS-Net, it was also decided for EARS-Vet not to monitor aminoglycosides and macrolides as antimicrobial classes, but as individual compounds, since they may have different resistance profiles (e.g. neomycin versus gentamicin or erythromycin versus tulathromycin).^3^ In the EFSA monitoring, antimicrobials are selected from a public health perspective and are monitored as individual compounds. Still, many of the antimicrobials monitored by EFSA in commensal *E. coli* have also been included in the EARS-Vet scope for *E. coli* (ampicillin, cefotaxime, chloramphenicol, nalidixic acid, ciprofloxacin, colistin, gentamicin, tetracycline, tigecycline, meropenem, imipenem and ertapenem).

Overall, the EARS-Vet scope would enable to make multiple comparisons between diseased animals, healthy food-producing animals and food thereof, and human patients. It would usefully complement the pool of data being analysed in the Joint Inter-Agency Antimicrobial Consumption and Resistance Analyses (JIACRA),^12^ to better understand the complex epidemiology of AMR across sectors and contribute to devising more efficient interventions against AMR. By covering major bacterial diseases in the main food-producing and companion animal species of Europe, EARS-Vet has the potential to make a difference in veterinary antimicrobial stewardship through the development of antimicrobial therapy guidelines, new AST interpretation criteria, or by revising indications of marketed antimicrobial products. Finally, it is noteworthy to point out that the EARS-Vet surveillance scope may evolve over time, as the epidemiology of AMR changes, more countries participate and the feasibility to monitor specific combinations changes.

## Acknowledgements

We are very grateful to all professionals involved in the definition of national surveillance scopes, consulted experts from the Federation of Veterinarians in Europe (FVE), ECDC, EMA, EFSA and EUCAST, as well as to Lucie Collineau (French Agency for Food, Environmental and Occupational Health and Safety) for providing sound feedback on this work. The views expressed in this publication are those of the authors and do not necessarily reflect the opinion of consulted experts and organisations.

## Transparency declaration

All authors have no conflicts of interest to disclose.

## References

1. World Health Organization. Global Action Plan on Antimicrobial Resistance. 2015. Available at: https://apps.who.int/iris/bitstream/handle/10665/193736/9789241509763_eng.pdf?sequence=1. Accessed January 7, 2021.

2. European Commission. A European One Health Action Plan against Antimicrobial Resistance. 2017. Available at: https://ec.europa.eu/health/sites/health/files/antimicrobial_resistance/docs/amr_2017_action-plan.pdf.

3. European Centre for Disease Prevention and Control. Antimicrobial resistance in the EU/EEA (EARS-Net) - Annual Epidemiological Report 2019. 2020. Available at: https://www.ecdc.europa.eu/sites/default/files/documents/surveillance-antimicrobial-resistance-Europe-2019.pdf. Accessed January 7, 2021.

4. European Centre for Disease Prevention and Control. EU protocol for harmonised monitoring of antimicrobial resistance in human Salmonella and Campylobacter isolates. 2016. Available at: https://www.ecdc.europa.eu/sites/portal/files/media/en/publications/Publications/antimicrobial-resistance-Salmonella-Campylobacter-harmonised-monitoring.pdf. Accessed October 25, 2020.

5. European Food Safety Authority, European Centre for Disease Prevention and Control. The European Union Summary Report on Antimicrobial Resistance in zoonotic and indicator bacteria from humans, animals and food in 2017/2018. 2020. Available at: file:///C:/Users/roudo/Desktop/EU-summary-report-antimicrobial-resistance-zoonotic-bacteria-humans-animals-2018.pdf. Accessed October 25, 2020.

6. Mader R, Damborg P, Amat J-P, et al. Building the European Antimicrobial Resistance Surveillance network in veterinary medicine (EARS-Vet). Euro Surveill 2021; 26.

7. World Organisation for Animal Health. Chapter 6.8: Harmonisation of National Antimicrobial Resistance Surveillance And Monitoring Programmes. In: Terrestrial Animal Health Code., 2018.

8. Hendriksen RS, Mevius DJ, Schroeter A, et al. Occurrence of antimicrobial resistance among bacterial pathogens and indicator bacteria in pigs in different European countries from year 2002 – 2004: the ARBAO-II study. Acta Veterinaria Scandinavica 2008; 50: 19.

9. Bourély C, Chauvin C, Jouy É, et al. Comparative epidemiology of E. coli resistance to third-generation cephalosporins in diseased food-producing animals. Veterinary Microbiology 2018; 223: 72–8.

10. World Health Organization (WHO). Global Antimicrobial Resistance Surveillance System: Manual for Early Implementation. 2015.

11. European Medicines Agency. Categorisation of antibiotics in the European Union. 2020. Available at: https://www.ema.europa.eu/en/documents/report/categorisation-antibiotics-european-union-answer-request-european-commission-updating-scientific_en.pdf. Accessed January 7, 2021.

12. European Centre for Disease Prevention and Control, European Food Safety Authority, European Medicines Agency. ECDC/EFSA/EMA second joint report on the integrated analysis of the consumption of antimicrobial agents and occurrence of antimicrobial resistance in bacteria from humans and food-producing animals – Joint Interagency Antimicrobial Consumption and Resistance Analysis (JIACRA) Report. EFSA Journal 2017; 15: 135.

